# LitRev: A data driven method for quick literature review from PubMed

**DOI:** 10.1101/2021.12.07.471694

**Authors:** Gourab Das

## Abstract

**Summary:** LitRev is a novel robust data driven approach, developed for quick literature review on a particular topic of interest. This method identifies common biological phrases that follow a power law distribution and important phrases which have the normalized point wise mutual information score greater than zero.

## 1 INTRODUCTION

Progressive advancement in the field of research and technology resulting exponential increment of scientific literatures in the repositories, manual extraction of required information from the articles has become a time consuming cumbersome task for researchers day by day. For many of them, it seems a difficult job to do a thorough literature review on a topic of interest at the very beginning of a research work within a time limit for a huge number of relevant article hits. Again, literature review is mandatory and unavoidable because this will show how far research has been progressed on a topic, what are the key ideas evolved, what are the techniques developed, what kind of tools become available, what can be the future aspects on that topic, and what the limitations are. When query returns a huge unmanageable number of hits, then, it is very troublesome and confusing to choose next query phrases for further in-depth search to retrieve relevant articles. In case of manageable number of hits also, some time is definitely needed for reading and writing the key points for further summarization and better understanding the available research work.

To address this problem for biological text data mining from Pubmed, variety methods including natural language processing (NLP), machine learning and rule based approaches for ranking articles, extracting topics and semantic-based keyword searching, have been successfully reviewed in [1]. Another efficient text mining java application, Biogyan (http://www.biogyan.com/) by Persistent LABS, enable us to do not only exhaustive combinatorial searching for a gene/protein, cell type/organelle, process or topic using databases like PubMed, NCBI Gene, UniProt, PDB, etc. but also fetching and ranking abstracts, highlighting relevant text, visualizing pathways and 3D structures have been facilitated on the basis of specialized scores. But major limitations of this tool are: a) Pubmed advanced queries can’t be used for searching, b) Can show only one abstract at a time and c) Can locate only certain sentence(s), gene or protein, cell type or organelle, process and query phrases in that abstract, hence overlook other important phrases often. Here, we develop a novel robust data driven method LitRev, for the identification of important key phrases from the Pubmed abstracts. This will provide a knowledgebase for current research and a source of future queries. Moreover, this method can fetch significant biological sequence patterns related to a particular gene/protein across species. See supplementary for results.

## 2 METHODS

To extract query related abstracts, a custom biopython script using NCBI E-utilities [2] is developed where Pubmed advanced queries can be used for fetching abstracts.

Pre-processing of the abstracts has been done by removing stop words [3], tags, numbers, punctuations, and multiple whitespaces. Stemming of the words has not been carried out in the preprocessing for efficient exact matching. Finally, all abstracts are converted into lowercase letters.

Phrases which are specific representation of words that appear frequently together, and infrequently in other contexts, has been found from the pre-processed abstracts using following formula [4]

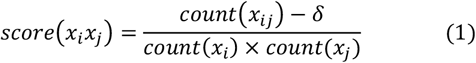

Document frequency (DF) is the count of abstracts where a single phrase or collection of phrases co-occurs at least once and calculated by another custom python script using NCBI E-utilities. The distribution of phrases in English language as well as in biomedical literatures follows a Power law type distribution partly [5-6]. A more suggested way to visualize such data is its complementary cumulative distribution function (CCDF) [7]. Power law fitting is performed and parameters (exponent α and cut-off x_min_) are estimated by maximum likelihood method. Derivation for maximum likelihood estimator for power law is in [8]. Goodness of fit is calculated using Kolmogorov-Smirnov (K-S) one sample test.

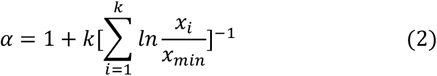

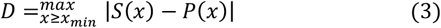

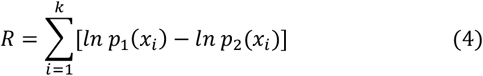

Then, data is fitted with several candidate distributions and compared to ensure the best fit distribution to data via likelihood ratio (R) test. Sign of log likelihood ratio (R) of the data under two competing distributions will decide which distribution fitted well to data. If it’s positive, data follows 1^st^ distribution else the 2^nd^ one. To make a quantitative judgment about the statistical significance of the observed sign of R, p-value is calculated using the method by Vuong et. al. [9]. If this p-value is small (say p < 0.05) then it is unlikely that the observed sign is a chance result of fluctuations and the sign is a reliable indicator of which model is the better fit to the data. On the other hand if p is large, the sign is not reliable and the test does not favour either model over the other. [7].

After locating commonly used biological phrases, ‘Normalized Point wise Mutual Information’ (NPMI) based correlation score [9] between query and remaining phrases is calculated to find desired important key phrases using following equation:

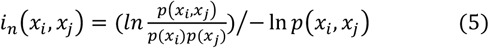

Phrase extraction is done by gensim and power law fitting, goodness fit test and comparison with other distributions is done by powerlaw package [10] under python.

## 3 RESULTS

Total number of phrases generated from **669** abstracts in response to query “clustered regularly interspaced short palindromic repeat AND hasabstract[text]” is **4480**. Due to data processing errors we had 38 phrases whose document frequencies are equal to zero. Eliminating them, we prepared out final dataset of **4442** phrases.

**Figure 1:**
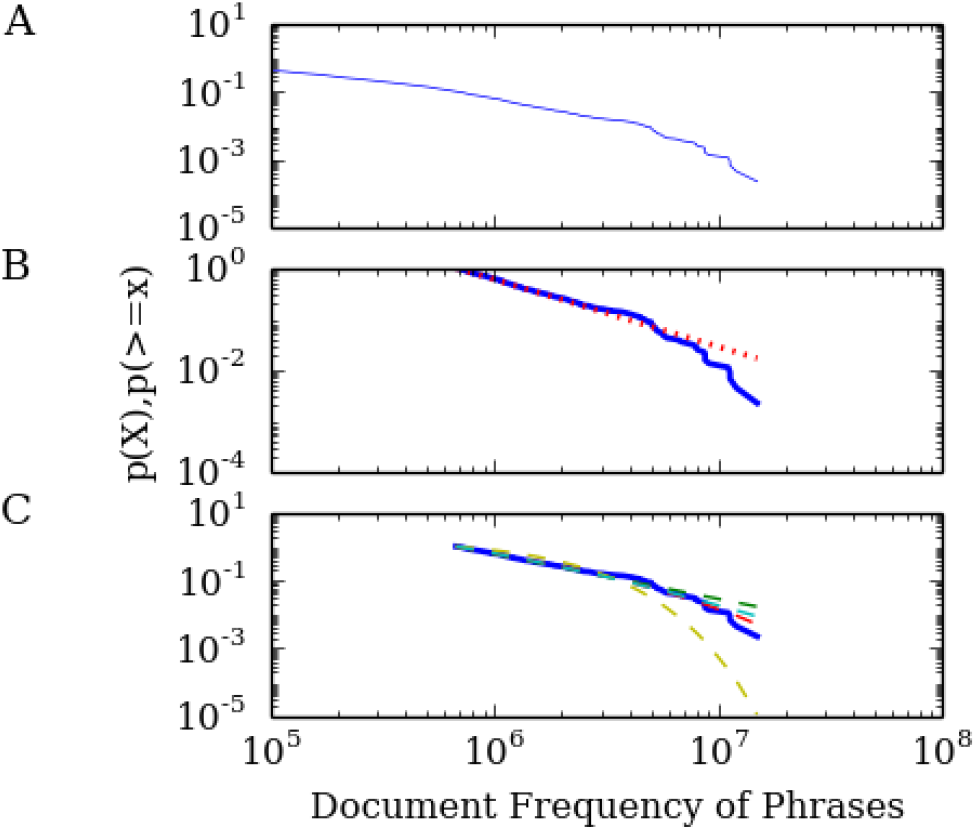
A. CCDF of phrases from Pubmed; B. Power law fitted in tail of the distribution; C. Comparison with other distributions. After plotting CCDF (Figure 1A) for this data and fitting (Figure 1B), estimated parameter values are **2.318** and **672983** for exponent α and cut-off (x_min)_ respectively, with KS distance **0.0332** and standard deviation 0.0621. So, power law is valid up to the data point having document frequency 672983. **450** data points which follow power law are the commonly used phrases with high document frequency. With the criteria of NPMI correlation score greater than some user defined cut-off (here, **0.1**), total **1578** phrases the important key phrases for the query. See supplementary for detailed list.

From Table 1, it implies significant sign of R (p< 0.05) that indicate truncated power law as the best fit.

**Table 1:**
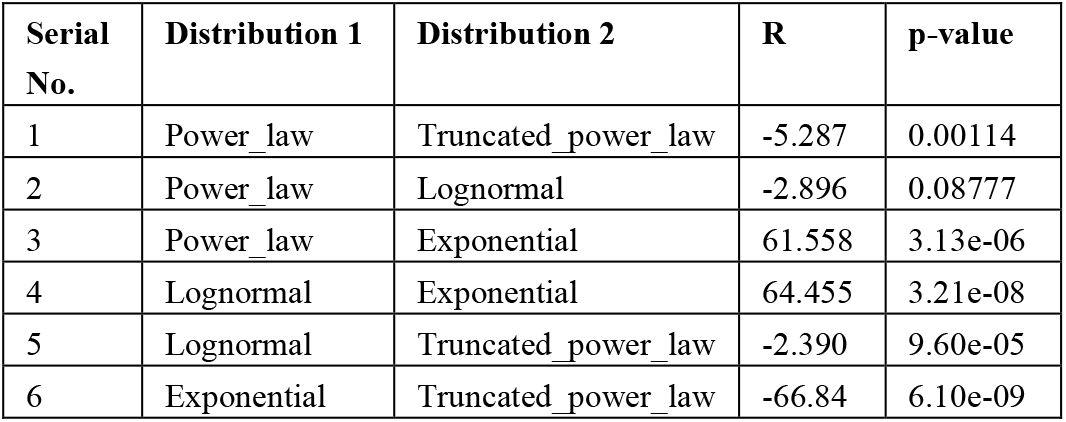
Comparison of fitting of data with different distributions

## 4 CONCLUSION

We successfully distinguish key phrases by studying and analyzing the truncated power law distribution of the phrases in Pubmed corpus and NPMI based correlation score between query and phrases. In spite of computational limitation for large number of hits, this data driven method can be a useful tool for literature review.

## ACKNOWLEDGEMENTS

I am sincerely thanking Prof. Indira Ghosh for reviewing and giving her valuable comments to enrich this manuscript.

